# Epigenetic centromere identity is precisely maintained through DNA replication but is uniquely specified among human cells

**DOI:** 10.1101/2022.09.07.506974

**Authors:** Megan A. Mahlke, Lior Lumerman, Peter Ly, Yael Nechemia-Arbely

## Abstract

Centromere identity is defined and maintained epigenetically by the presence of the histone variant CENP-A. How centromeric CENP-A position is specified and precisely maintained through DNA replication is not fully understood. The recently released Telomere-to-Telomere (T2T-CHM13) genome assembly containing the first complete human centromere sequences provides a new resource for examining CENP-A position. Mapping CENP-A position in clones of the same cell line to T2T-CHM13 identified highly similar CENP-A position following multiple cell divisions. In contrast, centromeric CENP-A epialleles were evident at several centromeres of different human cell lines, demonstrating the location of CENP-A enrichment and site of kinetochore recruitment varies among human cells. Across the cell cycle, CENP-A molecules deposited in G1 phase are maintained at their precise position through DNA replication. Thus, despite CENP-A dilution during DNA replication, CENP-A is precisely reloaded onto the same sequences within the daughter centromeres, maintaining unique centromere identity among human cells.

## Introduction

Centromere position is specified epigenetically by the centromeric histone H3 variant CENP-A (Barnhart et al., 2011; Black and Cleveland, 2011; Cleveland et al., 2003; Fachinetti et al., 2013). The precise deposition and maintenance of CENP-A across the cell cycle is the initial step in establishing functional centromeres, which are essential for assembling the kinetochore to allow for faithful chromosome segregation during mitosis, thereby preserving genomic integrity. CENP-A deposition in early G1 is tightly regulated by the combined activity of the Mis18 licensing complex (Fujita et al., 2007; Hayashi et al., 2004; Nardi et al., 2016; Pan et al., 2017; Spiller et al., 2017; Stellfox et al., 2016) and CENP-A’s chaperone HJURP (Barnhart et al., 2011; Dunleavy et al., 2009; Foltz et al., 2009; Pan et al., 2019; Tachiwana et al., 2015; Wang et al., 2014; Zasadzińska et al., 2013). During subsequent DNA replication, HJURP (Zasadzińska et al., 2013), CENP-C and the CCAN (Nechemia-Arbely et al., 2019) are required for the retention and reloading of CENP-A at centromeres of daughter DNA strands. CENP-A distribution between the two daughter centromeres during DNA replication results in dilution of CENP-A to ∼50% occupancy until the following G1 (Jansen et al., 2007; Nechemia-Arbely et al., 2012; Silva et al., 2012; Stankovic et al., 2017). The temporal separation between CENP-A dilution in S phase and new CENP-A deposition in G1 raises the important question of how centromere epigenetic identity is maintained across the cell cycle (Mahlke and Nechemia-Arbely, 2020).

Human centromeres are megabase-long chromosomal regions (Wevrick and Willard, 1989) composed of tandemly repeated 171 base pair (bp) α-satellite monomers organized into High-Order Repeat (HOR) arrays (Willard, 1985; Willard and Waye, 1987) that cannot be resolved with traditional sequencing approaches (Eichler et al., 2004; Miga, 2015). Thus, centromeres in the hg38 human genome assembly (GRCh38p.13) are represented by models that reflect the observed variation in human α-satellite repeat sequences but do not distinguish between centromeric and pericentromeric sequences, and arbitrarily assign the order of repeats in each centromeric array (Levy et al., 2007; Miga et al., 2014; Schneider et al., 2017). Moreover, attempts to use sequencing-based approaches to study centromere epigenetics are complicated by the inability to accurately map short sequencing reads to a precise centromeric location or to interpret whether mapped sequencing reads represent the true position of epigenetic modifications in centromeric chromatin. Consequently, the highly complex repetitive nature of human centromere sequences hinders the study of centromere genomics and epigenetics, limiting our understanding of CENP-A maintenance.

The complete CHM13 human genome assembly released by the Telomere-to-Telomere (T2T) Consortium represents a significant achievement that uncovers the true sequences of large and repetitive genomic elements using long-read sequencing technologies (Nurk et al., 2022), including 156.2 Mb of centromeric satellites that remained missing from the hg38 assembly (GRCh38p.13) 21 years after the first human genome was released (Lander et al., 2001). Here, we used the T2T assembly (CHM13v1.1) that contains the first accurate description of human centromere sequences to assess the pattern of centromeric CENP-A binding in various human cell types and across the cell cycle. Using new CENP-A ChIP-seq datasets, as well as our own (Nechemia-Arbely et al., 2019) and other publicly available CENP-A ChIP-seq and CUT&RUN data (Dumont et al., 2020; Logsdon et al., 2021; Thakur and Henikoff, 2016), we demonstrate that CENP-A position is highly similar in a single cell line and its derived clones following multiple cell divisions, but varies significantly between different human cell lines, indicating the presence of CENP-A epialleles at several human centromeres. Further, we test the sequences bound by CENP-A in G1 and G2 enriched cells and demonstrate that CENP-A is maintained at the same α-satellite sequences by precise reloading at the same position during DNA replication, thereby preserving centromere epigenetic identity.

## Results

### The position and distribution of CENP-A at human centromeres differs significantly between hg38 and CHM13 T2T genome assemblies

Though the position of α-satellite monomer repeats in the centromere models contained within the hg38 assembly (GRCh38p.13) are arbitrary and do not represent true centromere HOR arrangement, the hg38 centromere models have been used to estimate the distribution of CENP-A at human centromeres (Hoffmann et al., 2020; Nechemia-Arbely et al., 2019). To investigate the true position of CENP-A at human centromeres using the recently released T2T genome assembly (CHM13v1.1) (Nurk et al., 2022), we performed CENP-A ChIP-seq in PD-NC4 fibroblasts (Amor et al., 2004). To obtain sufficient number of cells for CENP-A ChIP and to derive single-cell clones, we first immortalized and partially transformed PD-NC4 cells by expressing *hTERT* and oncogenic *KRAS*^*V12*^, respectively. We mapped the resulting data (Supplementary Table 1) alongside previously generated CENP-A ChIP-seq data from HeLa cells (Nechemia-Arbely et al., 2019) to the T2T (T2T-CHM13v1.1) and hg38 (GRCh38p.13) genome assemblies, retaining all high-quality centromeric reads and assigning multi-mapping repetitive reads randomly to a single location (Fig. 1A, top panel).

**Figure 1.**
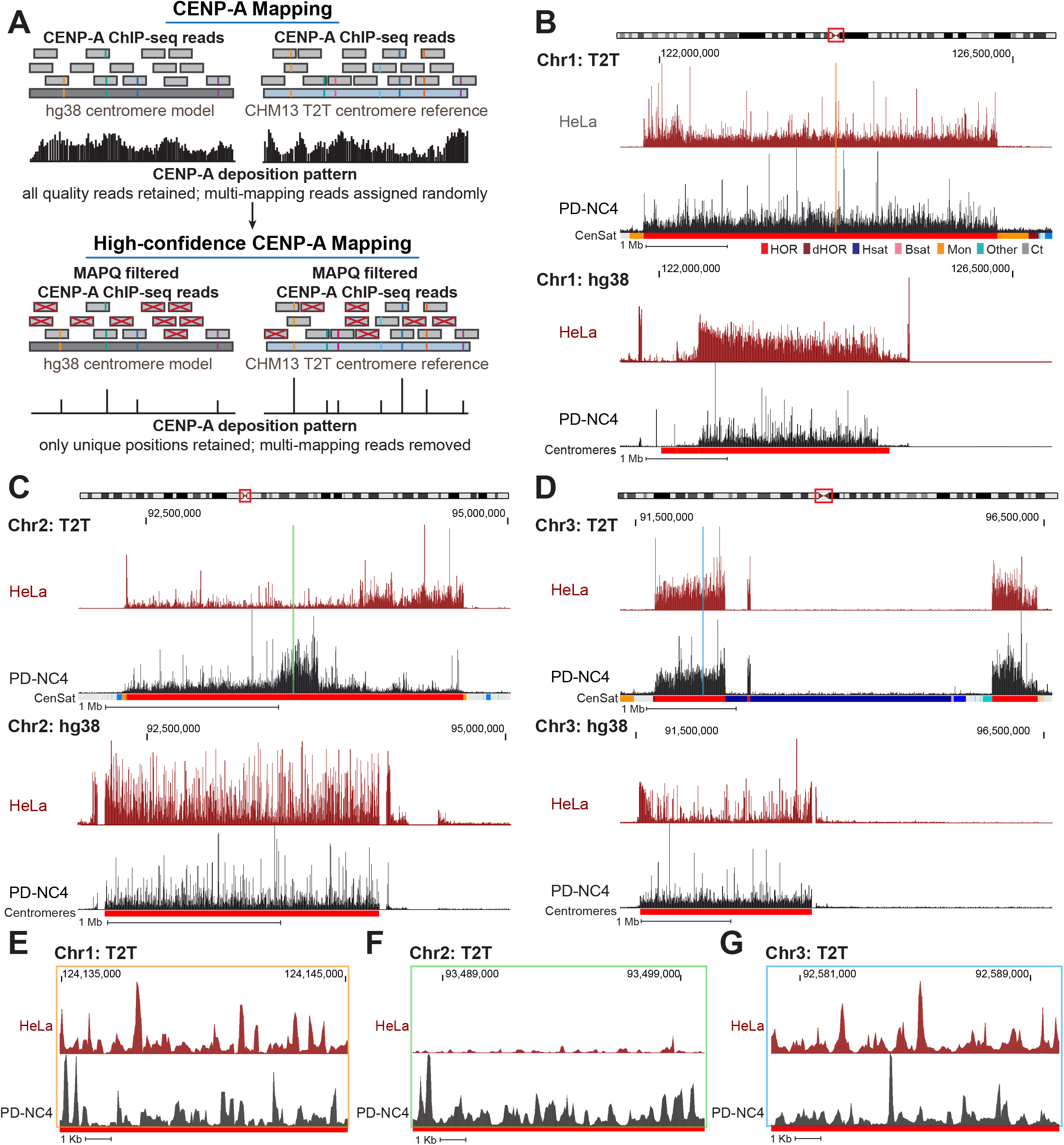
CENP-A alignment patterns differ dramatically between hg38 centromere reference models and T2T centromere sequences. **(A)** Schematic representing conventional mapping of CENP-A-bound reads (top panel) and MAPQ filtered high-confidence mapping of CENP-A-bound reads (bottom panel) to the hg38 (left) and T2T (right) assemblies. The higher number of unique variants in the T2T assembly allows for a greater number of high-confidence CENP-A reads to be mapped. Reads with colored lines represent unique variants within the α-satellite repeats. **(B-D)** Position of CENP-A reads in HeLa (maroon) and PD-NC4 (black) cells when mapped to the T2T (top) and hg38 (bottom) assemblies at the centromeres of chromosome 1 (**B**), 2 (**C**), and 3 (**D**). Chromosome ideograms at top indicate position of centromere. Annotation tracks below the T2T assembly indicate positions of centromeric satellites; see color key in panel **B**. (HOR=higher order repeats, red; dHOR=divergent higher order repeats, maroon; Hsat=human satellites, blue; Bsat=Beta satellites, pink; Mon=monomers, orange; Others= Other centromeric satellites, teal; Ct= Centric Satellite Transition Regions, grey). Annotation tracks below the hg38 assembly indicate positions of centromeres (red). Scale bar, 1 Mb. **(E-G)** High-resolution view of CENP-A reads in HeLa (maroon) and PD-NC4 (black) cells when mapped to the T2T assembly at the same centromeres shown in B-D. Color coded panels in E-G represent the colored locations indicated in (**B**-**D**). Scale bar, 1 Kb.

Mapping of CENP-A-bound DNA reads revealed dramatic differences in CENP-A position at nearly all centromeres between the hg38 and T2T genome assemblies (Figure 1B-D, Supplementary Fig. 1). Some differences in CENP-A mapping between the two assemblies were the result of uncovering the actual arrangement of HORs and other repetitive elements at centromeres in the T2T assembly. At chromosomes 1 (Fig. 1B), 6, 19 and 22 (Supplementary Fig. 1), the T2T assembly contains a longer span of centromeric HOR sequences than was predicted in the hg38 centromere model. At chromosomes 3 (Fig. 1D) and 4 (Supplementary Fig. 1), the T2T assembly includes large insertions of human satellite (Hsat) sequences in the center of the HOR array (Altemose et al., 2022) splitting CENP-A into two separate centromeric regions and dramatically changing the centromeric CENP-A landscape between the hg38 and T2T assemblies. At other centromeres, we discovered distinctive CENP-A enrichment patterns between the two assemblies that were not clearly related to changes in HOR size or organization. At the centromere of chromosome 2, CENP-A is distributed uniformly across the HOR in both HeLa and PD-NC4 cells aligned to the hg38 assembly but is locally enriched in distinct centromeric regions in HeLa and PD-NC4 cells when aligned to the T2T assembly (Fig. 1C). These cell line-specific regions of CENP-A enrichment were also evident at the centromeres of other chromosomes (See chromosomes 6, 8, 9, 11, 17, 18, 22, and X in Supplementary Fig. 1). Alignment to the T2T assembly also highlighted the small-scale differences in CENP-A position between HeLa and PD-NC4 cells (Fig. 1E-G). These striking differences in CENP-A position between HeLa and PD-NC4 cells aligned to the T2T assembly suggest that the centromeric site enriched for CENP-A binding may be specified at different positions within the same centromeres in different human cells.

### CENP-A position varies between human cell lines

Recent reports have identified that human centromeres evolve through layered expansions leading to generation of distinct sets of α-satellite repeats within the same HOR that can be clustered based on their shared sequence variants into higher order repeat haplotypes or “HOR-haps” (Altemose et al., 2022). Furthermore, CENP-A enrichment and site of kinetochore recruitment in CHM13 cells was reported to frequently occur at the evolutionarily younger HOR-hap and site of recent expansion (Altemose et al., 2022). Despite this progress in understanding centromere evolution and its effect on centromere structure, the location of centromeric CENP-A enrichment at all centromeres among diverse human cells remains largely unexplored.

By mapping CENP-A ChIP-sequencing data from CHM13, PD-NC4 and HeLa cells to the T2T assembly and using the recent characterization of the active HOR S3CXH1L (DXZ1 in hg38) within the X centromere into distinct HOR-haps (Altemose et al., 2022), we found that CENP-A in PD-NC4 and CHM13 cells is enriched at the evolutionarily younger HOR-hap and site of recent HOR expansion within the S3CXH1L HOR, while CENP-A in HeLa cells is enriched at an evolutionarily older HOR-hap of S3CXH1L that does not overlap a site of recent expansion (Fig. 2A). In contrast, examination of the active HOR of chromosome 2, S2C2H1L (D2Z1 in hg38), revealed that CENP-A in both CHM13 and HeLa cells is enriched at an evolutionarily older HOR-hap and site of recent HOR expansion, while in PD-NC4 cells, CENP-A is enriched at an evolutionarily younger HOR-hap that also overlaps a site of recent expansion (Fig. 2B). Previous work has shown that two α-satellite HOR arrays within the centromere of chromosome 17, S317H1L (D17Z1) and S317H1-B (D17Z1-B), are both capable of recruiting CENP-A and forming a kinetochore in GM08828 (lymphocyte) cells (Maloney et al., 2012). In the cell lines we examined, CENP-A was only found at the S317H1L (D17Z1) HOR but was localized at different HOR-haps within the array (Fig. 2C). In CHM13 cells, CENP-A is enriched at the evolutionarily younger HOR-hap and is highly enriched at the zone of recent expansion. In PD-NC4 cells, CENP-A is enriched at an evolutionarily older HOR-hap flanking the right side of the younger HOR-hap (at T2T coordinates of ∼27 Mb). In HeLa cells, CENP-A is enriched at an evolutionarily older HOR-hap that flanks the opposite side of the younger HOR-hap (at T2T coordinates of ∼24 Mb). Interestingly, HeLa cells also exhibit a lower level of CENP-A enrichment at the evolutionarily older HOR-hap flanking the right side of the younger HOR-hap (at T2T coordinates of ∼27 Mb), which may indicate that HeLa cells have distinct maternal and paternal CENP-A epialleles at the centromere of chromosome 17 (Fig. 2C). Taken together, these distinct patterns of centromeric CENP-A enrichment in different human cell lines suggest the presence of previously undocumented CENP-A epialleles (i.e., different positioning of the region enriched for CENP-A binding with respect to the underlying α-satellite DNA) within a single human centromeric HOR.

**Figure 2.**
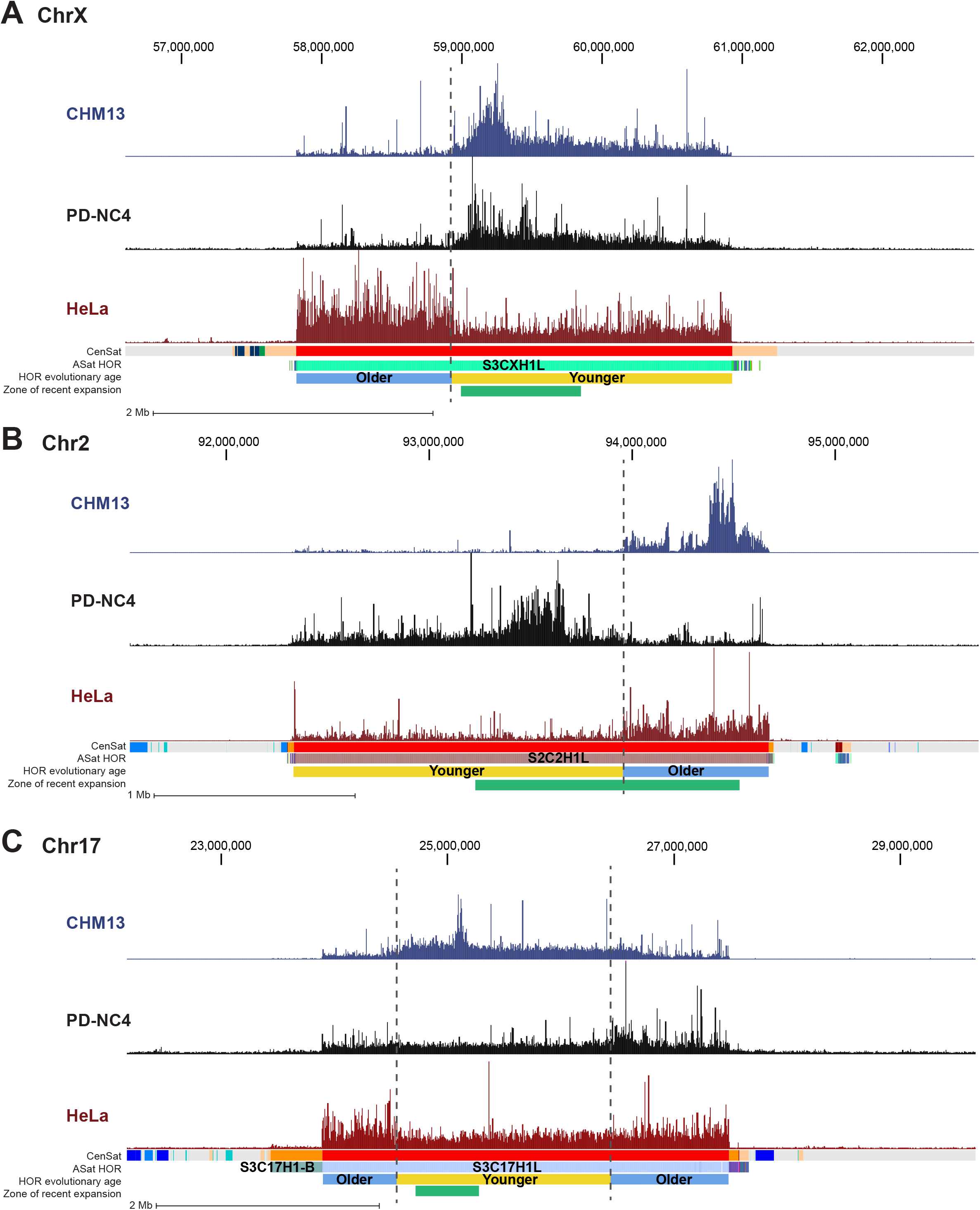
CENP-A binds at evolutionarily younger and older HOR-haps in different centromeres and different cell lines. **(A-C)** Mapping of CENP-A-bound reads in CHM13 (blue), PD-NC4 (black), and HeLa cells (maroon) at the centromeres of chromosome X (**A**), chromosome 2 (**B**), and chromosome 17 (**C**). Dashed lines mark regions that are differentially enriched for CENP-A binding between cell lines. Annotation tracks indicate positions of centromeric satellites (CenSat), higher order repeat arrays (ASat HOR), rough HOR age based on evolutionary age of HOR-haps (HOR evolutionary age), and the location of the zone of recent HOR expansion (Zone of recent expansion). HOR age and Zone of recent expansion were adapted from (Altemose et al., 2022).

To further investigate centromeric CENP-A epialleles in human cell lines, we compared CENP-A position in HeLa (cervical cancer), PD-NC4 (fibroblast), RPE-1 (retinal pigment epithelium) and HuRef (lymphoblastoid cell line (Levy et al., 2007)) cells when aligned to the T2T assembly, using new (PD-NC4) and previously published CENP-A ChIP-seq [(Nechemia-Arbely et al., 2019) for HeLa; (Logsdon et al., 2021) for CHM13; (Henikoff et al., 2015) for HuRef)] and CUT&RUN [(Dumont et al., 2020) for RPE-1] datasets. We found that CENP-A position varies between human cell lines in the centromeres of almost all chromosomes (excluding chromosomes 4, 15, 16, and 21, at which CENP-A pattern is similar across cell lines) when mapped to the T2T genome assembly (Fig. 3A-C, Supplementary Figs. 2 and 3). Conversely, positions of centromeric CENP-A are highly similar in PD-NC4 parental cells and two PD-NC4 single-cell derived clones following multiple cell divisions (Fig. 3G), suggesting that the observed CENP-A epialleles between human cell lines of different origin truly represent individual or cell type-specific changes in CENP-A position and that within the same cell type CENP-A position is maintained over many cellular divisions. At some centromeres, a single region enriched for CENP-A binding was easily identified, although the location of the region enriched for CENP-A may differ between cell lines (Fig. 2A-B; Fig. 3A,C; Supplementary Figs. 2 and 3 – See centromeres of chromosomes 2, 6, 8, 10, 12, 14, 16, 17, 18, 20). At other centromeres, two regions enriched for CENP-A binding were evident, suggesting differential CENP-A position between maternal and paternal centromeres/chromosomes in diploid cell lines (Fig. 3A RPE-1 cells; Fig. 3C PD-NC4 cells). However, a definitive region enriched for CENP-A was challenging to identify in some centromeres of different cell lines (Fig. 3B; Supplementary Figs. 2 and 3, for example see the HeLa and PD-NC4 centromeres of chromosomes 3 and 4, as well as all lines in chromosome 21).

**Figure 3.**
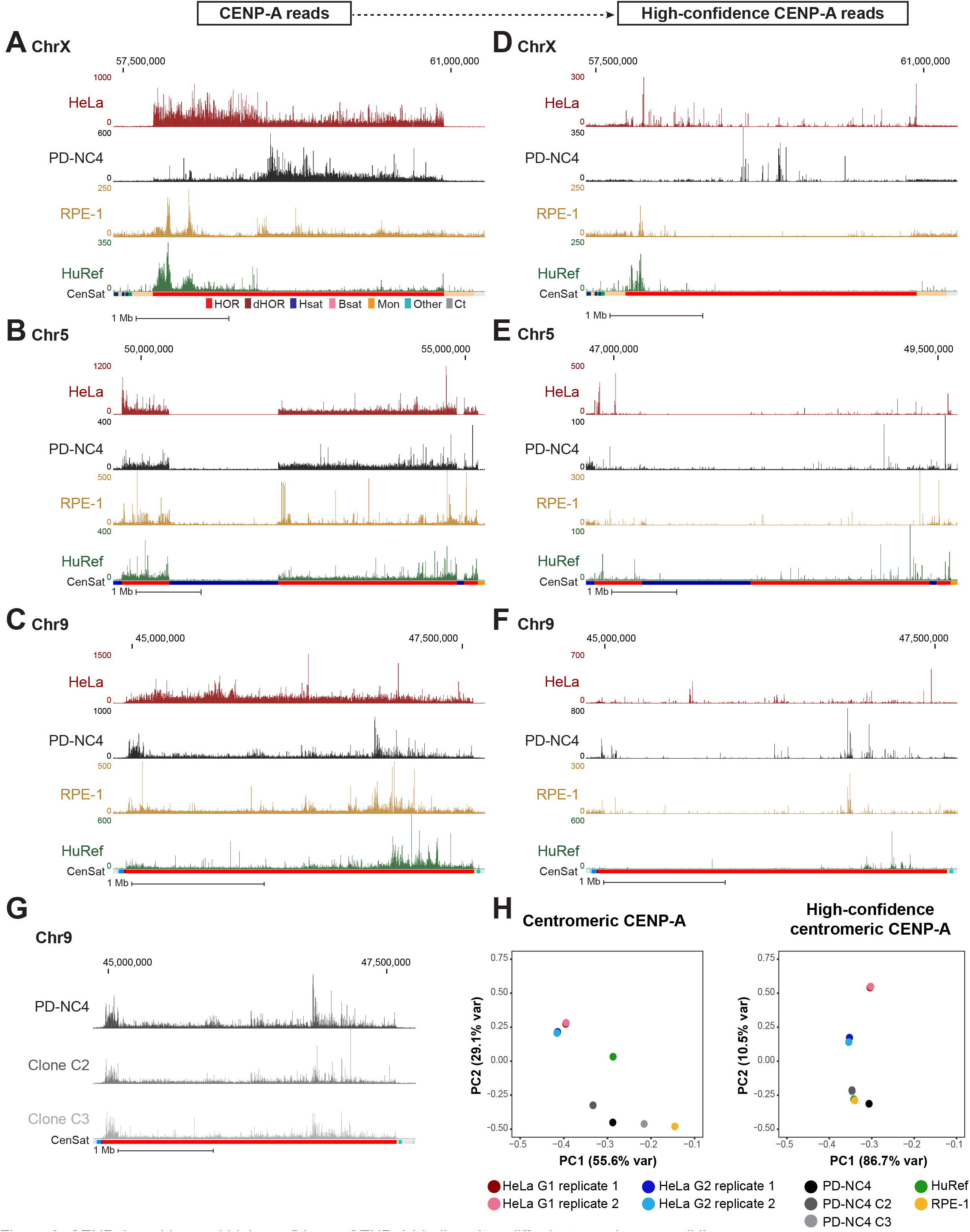
CENP-A position and high-confidence CENP-A binding sites differ between human cell lines. **(A-C)** Mapping of CENP-A to chromosome X (**A**), chromosome 5 (**B**), and chromosome 9 (**C**) of the T2T assembly in HeLa (maroon), PD-NC4 (black), RPE-1 (yellow), and HuRef (green) cells. Annotation tracks below the T2T assembly indicate positions of centromeric satellites; see color key in panel **A**. (HOR=higher order repeats, red; dHOR=divergent higher order repeats, maroon; Hsat=human satellites, blue; Bsat=Beta satellites, pink; Mon=monomers, orange; Others= Other centromeric satellites, teal; Ct= Centric Satellite Transition Regions, grey). Scale bar, 1 Mb. **(D-F)** Mapping of high-confidence CENP-A-bound reads to chromosome X (**D**), chromosome 5 (**E**), and chromosome 9 (**F**) of the T2T assembly in HeLa, PD-NC4, RPE-1, and HuRef cells. Scale bar, 1 Mb. **(G)** Mapping of CENP-A-bound reads from parental PD-NC4 fibroblasts and two single cell-derived clones to the centromere of chromosome 9 in the T2T assembly. **(H)** PCA plots of significantly enriched centromeric CENP-A peaks (MACS2, p< 0.00001) at all (left panel) and high-confidence (right panel) positions in different human cell lines. HeLa G1 replicates, maroon and pink; HeLa G2 replicates, dark and light blue; PD-NC4 parental and derived clones, black and grey; RPE1, yellow; HuRef, green.

Recognizing that conventional mapping of CENP-A-bound DNAs can generate a bias due to random assignment of multi-mapping reads, we produced high-confidence centromeric CENP-A alignments, which can be achieved with two methods: 1) Filtering each dataset to retain only high-confidence CENP-A alignments based on MAPQ score (MAPQ>20), effectively removing any reads that map to more than one location and retaining only the highest confidence alignments at unique centromeric sequences (Fig. 1A, bottom panel), or 2) K-mer assisted mapping that identifies unique k-mers within centromeric HORs and retains reads mapping to these locations only (Altemose et al., 2022). Mapping of high-confidence CENP-A reads using MAPQ and k-mer approaches were highly concordant (Supplementary Fig. 4). We therefore chose to continue with the MAPQ approach, which is consistent with our previous methodology (Nechemia-Arbely et al., 2019). Though retaining only the highest confidence reads prevents the obfuscation of CENP-A position due to random mapping, it also removes approximately 90-95% of centromeric CENP-A reads due to the highly repetitive nature of centromeric HOR arrays, limiting the analysis to only 5-10% of CENP-A-bound DNAs (Fig. 1A; Supplementary Table 2). Of note, the RPE-1 CUT&RUN dataset retained a much higher percentage (24%) of reads after MAPQ filtering compared to HeLa, PD-NC4 and HuRef ChIP-seq datasets (Supplementary Table 2). We found that high-confidence CENP-A positions also differ between human cell lines at most centromeres (Fig. 3D-F; Supplementary Figs. 2-3), supporting our findings using conventional high-quality CENP-A mapping and validating the existence of previously undocumented CENP-A epialleles within a single centromeric HOR at several human centromeres. However, retaining only high-confidence CENP-A reads reduced the apparent variability in CENP-A deposition patterns between human cell lines, as evidenced by PCA analysis (Fig. 3H), highlighting the importance of both conventional high-quality mapping and high-confidence mapping strategies when evaluating CENP-A position and distribution at centromeres.

### CENP-A position is maintained through DNA replication with precision

Using CENP-A ChIP-sequencing from HeLa cells enriched in G1 and G2 and mapped to the centromere reference models within the hg38 assembly, we previously showed that CENP-A is precisely retained at the same centromeric sequences through DNA replication (Nechemia-Arbely et al., 2019). Since our current analysis reveals that mapping of the same data to both hg38 and T2T genome assemblies yields significantly different CENP-A distribution patterns (Fig. 1B-G), we further tested our hypothesis of precise CENP-A retention through DNA replication by mapping our previous CENP-A ChIP-sequencing data from HeLa cells enriched in G1 and G2 (Nechemia-Arbely et al., 2019) to the T2T genome assembly. Centromeric CENP-A distribution in cells enriched in G1 phase was highly similar to its distribution in cells enriched in G2 phase at all human centromeres (Fig. 4A-B, E-F; Supplementary Fig. 5). Almost all (∼93%) significantly enriched CENP-A peaks at α-satellite sequences (MACS2 p<0.00001, ≥ 10-fold enrichment; see methods) present during G1 remained at the same sequences in G2 (Fig. 4I), demonstrating CENP-A retention at the same centromeric sequences through DNA replication.

**Figure 4.**
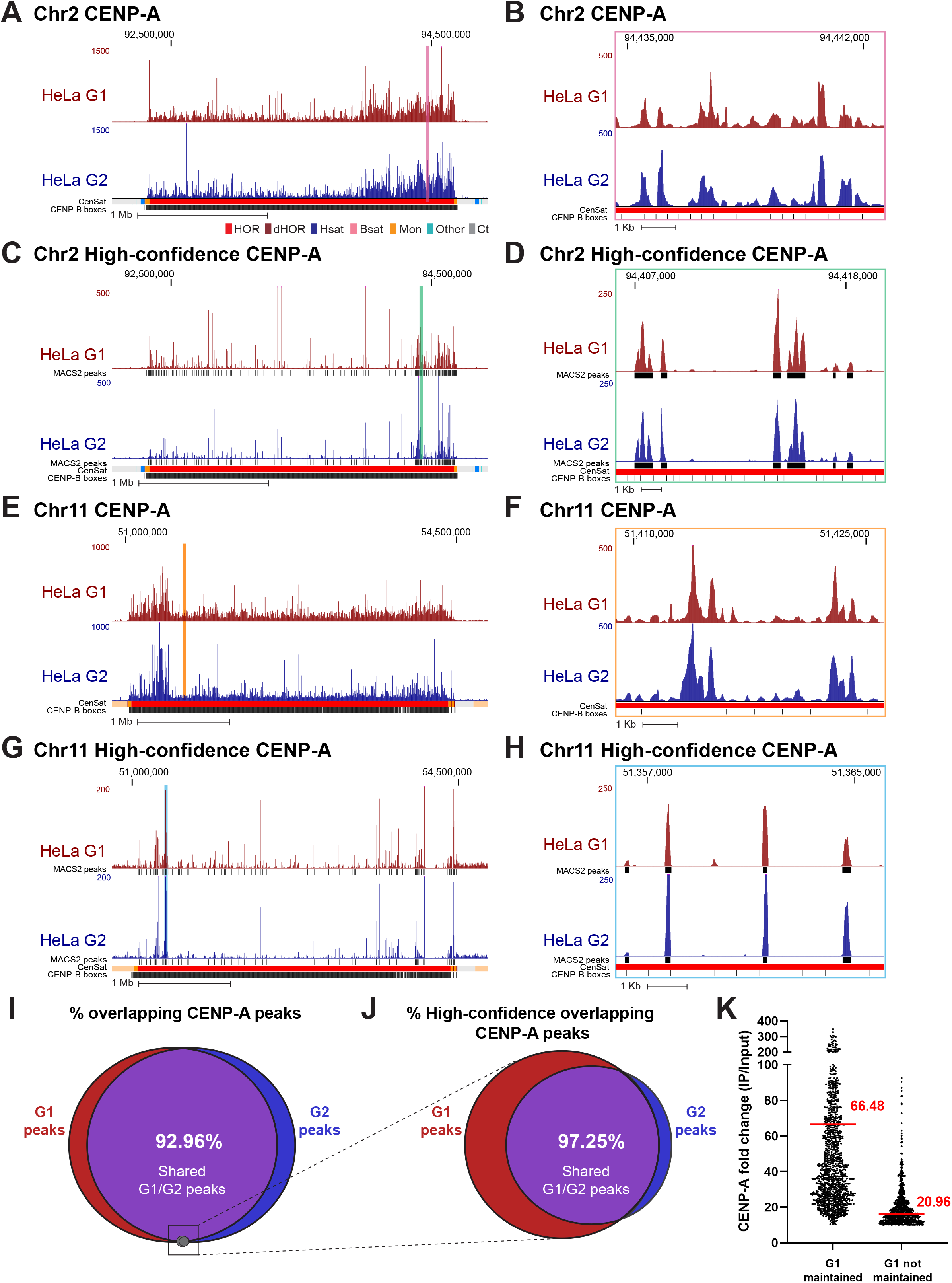
CENP-A nucleosomes are retained through DNA replication at human centromeres. **(A-D)** CENP-A position in HeLa cells enriched at G1 (maroon) and G2 (blue) mapped to chromosome 2 of the T2T assembly with conventional mapping (**A-B**) and high-confidence mapping (**C-D**). CENP-A peaks are highly concordant between HeLa G1 and G2 data at smaller (**A, C -** scale bar 1 Mb) and higher-resolution (**B, D -** scale bar 1 Kb) views. Color coded panels (**B, D**) represent high resolution view of the colored locations indicated in **A, C**. MACS2 peaks in **C** and **D** in black indicate positions of significantly enriched for CENP-A (MACS2 p<0.00001). **(E-H)** CENP-A position in HeLa cells enriched in G1 and G2 and mapped to chromosome 11 of the T2T assembly, with conventional mapping (**E-F**) and high-confidence mapping (**G-H**). CENP-A peaks are highly concordant between HeLa G1 and G2 data at smaller (**E, G -** scale bar 1 Mb) and higher-resolution (**F, H-** scale bar 1 Kb) views. Color coded panels (**F, H**) represent high-resolution view of the colored locations indicated in (**E, G**). MACS2 peaks in **G** and **H** indicate positions significantly enriched for CENP-A (MACS2 p<0.00001). Annotation tracks indicate positions of centromeric satellites (CenSat) and position of CENP-B boxes. **(I)** Venn diagram depicting the percent overlap of significantly enriched CENP-A peaks identified in data from HeLa cells enriched in G1 and G2 phase (MACS2 p<0.00001, ≥ 10-fold enrichment). 93% of CENP-A peaks in G1 were retained in G2. Small grey circle represents the relative size (5.1%) of the high-confidence CENP-A peaks depicted in the venn diagram in **J. (J)** Venn diagram depicting the percent overlap of significantly enriched CENP-A peaks identified in high-confidence mapped reads from HeLa cells in G1 and G2 phase (MACS2 p<0.00001, ≥ 10-fold enrichment). 97% of CENP-A peaks in G2 were precisely maintained from their existing position in G1. (**K**) Enrichment levels of significantly enriched (p<0.00001, ≥ 10-fold enrichment) high-confidence CENP-A G1 peaks that are either maintained or not maintained into G2. Peaks that were maintained from G1 to G2 had an average fold enrichment value of 66.48. Peaks that were not maintained from G1 to G2 had an average fold enrichment value of 20.96.

Across all centromeres in the hg38 assembly, high-confidence mapping previously identified 96 unique single-copy CENP-A binding sites in HeLa cells (Nechemia-Arbely et al., 2019). Using the same HeLa dataset aligned to the T2T assembly, we identified 1,879 unique single-copy CENP-A binding sites in HeLa cells that are significantly enriched for CENP-A (MACS2 p<0.00001, ≥ 10-fold enrichment). Within this 20-fold expanded set of unique high-confidence CENP-A binding sites, ∼97% of G2 CENP-A peaks were maintained at their G1 positions, demonstrating precise reloading of CENP-A at the same position during DNA replication (Fig. 4C, G, J). High-resolution nucleosomal views demonstrate retention of CENP-A nucleosomes through DNA replication at the same sequence and centromeric position with precision (Fig. 4D, H, Fig. S5B). 33.7% of high-confidence CENP-A G1 peaks were not maintained from G1 to G2 (Fig. 4J, red). However, these high-confidence lost G1 peaks were ∼3.2x-fold less enriched on average for CENP-A binding compared to G1 peaks that were maintained in G2 (Fig. 4K, Fig. S5C), indicating that lost G1 peaks were not a hot spot for CENP-A deposition at G1 within the cell population and represent sites that are less likely to be occupied by CENP-A. Moreover, since the MAPQ filtered dataset represents only 5.63% of total CENP-A-bound DNAs in HeLa (Supplementary Table 2) and 5.1% of significantly enriched CENP-A peaks (Fig. 4I, grey circle), the lost G1 peaks represent only 1.89% of total CENP-A-bound DNAs in G1, while most G1 peaks (∼93%) are retained through DNA replication (Fig. 4I). Taken together, the overwhelming majority of centromeric CENP-A enriched peaks were maintained at the same sequences and at the same position through DNA replication, demonstrating precise retention of centromeric CENP-A nucleosomes during DNA replication.

## Discussion

The first assembly of complete human centromere sequences as part of an entire human genome (T2T-CHM13) increases the accuracy of mapping CENP-A nucleosomes within these highly complex genomic loci. While the true centromere sequences remain highly challenging for short read mapping approaches, they vastly increase the number of known unique centromeric variants that can be used to confidently anchor the positions of centromeric proteins (Aganezov et al., 2022).

Using conventional and high-confidence mapping approaches we demonstrate that CENP-A position at centromeres varies significantly between human cell lines mapped to the T2T assembly (Figs. 1-3). This indicates that different human cell lines have distinct CENP-A epialleles at several centromeres and that the site of CENP-A enrichment, which recruits the kinetochore, is found at different locations within centromeric α-satellite DNA. Indeed, CENP-A epialleles have been documented on different HORs within the centromere of human chromosome 17 (Maloney et al., 2012) and at different locations within a single HOR in human chromosome X (Altemose et al., 2022). CENP-A epialleles have also been documented in inbred mouse strains (Arora et al., 2022). Here, we find that CENP-A epialleles occur at different locations within the same HOR array of a centromere and may correlate with distinct centromeric HOR-haps. In contrast, within the same cell line, centromeric CENP-A position is maintained from parental cells to single-cell derived clones, demonstrating faithful CENP-A maintenance across many cell divisions. The underlying cause for the differences in CENP-A position we observed in this diverse panel of human cell types remains to be determined.

One explanation for differences in CENP-A position could be innate genomic variability in centromere HOR array size and/or organization at the individual and population level (Miga, 2019) which is likely to differ between human cell lines of distinct origins. Thus, enrichment of CENP-A at different positions in human cell lines could reflect over-representation of a portion of CENP-A-bound reads in a sample due to genomic expansion of a particular HOR, or HOR-hap, relative to the CHM13-T2T assembly. CENP-A position at centromeres can also differ between maternal and paternal chromosomes in normal diploid cell lines (Maloney et al., 2012; Miga, 2019), which may further complicate our interpretation of CENP-A position when aligned to the functionally haploid CHM13-derived T2T centromere sequences (Altemose et al., 2022). Alternatively, the possibility that CENP-A position at centromeres may change during development of different cell types (fibroblast, lymphocyte, etc.) within an individual has yet to be investigated and could also contribute to the distinct localization of CENP-A within this diverse panel of human cell types.

Our results also reveal that it can be difficult to identify a single region of CENP-A enrichment within the centromere HOR when aligning non-CHM13 cell lines to the T2T assembly in many of the centromeres (Supplementary Fig. 1). Previous studies of CENP-A position using the T2T assembly have primarily used the CHM13 cell line, as this approach minimizes complications that may arise from mapping data generated from other human cell lines to the CHM13-derived T2T assembly. However, until individualized telomere-to-telomere reference genomes are produced for multiple human cell lines, the CHM13-derived T2T assembly is the best tool for mapping to centromeres and other previously unresolved repetitive regions. The extent to which our observations reflect true differences in CENP-A enrichment patterns between human cell lines or individuals rather than differences in centromere HOR composition or maternal/paternal effects in diploid human cell lines is an extremely interesting question that can only be resolved by generating centromere assemblies for several individual human cell lines.

The T2T assembly has allowed us to identify a 20-fold increase (from 96 to 1,879) in the number of unique centromeric variants at which CENP-A is bound (Aganezov et al., 2022). Using this expanded set of unique centromeric markers, we find that CENP-A position is precisely maintained through DNA replication by reloading onto the exact same α-satellite sequence and position during DNA replication (Fig. 4), consistent with our previous findings using the human centromere models in the hg38 assembly. This supports our proposed model for the epigenetic maintenance of human centromeres by precise CENP-A reloading and selective CENP-A retention at centromeres coupled with removal of ectopic CENP-A during DNA replication (Nechemia-Arbely et al., 2019). Collectively, our findings demonstrate plasticity in the position of CENP-A enrichment and location of kinetochore recruitment between human cells, accompanied by precise CENP-A maintenance across the cell cycle to preserve epigenetic centromere identity.

## Methods

### Cell culture

PD-NC4 fibroblasts (Amor et al., 2004) (a kind gift from Ben Black) were immortalized by ectopic retroviral expression of human telomerase (*hTERT*) and oncogenic *KRAS*^*V12*^. Retroviral supernatants were obtained from 293GP cells co-transfected with pVSV-G and pBABE-puro-hTERT or pBABE-hygro-KRAS-V12 constructs (kind gifts from Jerry Shay) using FuGENE HD (Promega) and passed through a 0.45 um filter. Following transduction of PD-NC4 cells in the presence of 5 ug/mL polybrene (Santa Cruz), infected cells were selected with 2 ug/mL puromycin and 100 ug/mL hygromycin, respectively. Cells were maintained in DMEM medium (Gibco) containing 10% fetal bovine serum (Omega Scientific), 100U/ml penicillin, 100U/ml streptomycin and 2mM l-glutamine at 37oC in a 5% CO2 atmosphere with 21% oxygen. Cells were maintained and split every 3-4 days according to ATCC recommendations. Single cell-derived clones were obtained by limited dilution.

### Chromatin extraction and purification

Chromatin was extracted Nuclei from 1×10^8^ nuclei of PD-NC4 cells as previously described (Nechemia-Arbely et al., 2019). CENP-A-bound chromatin was immunoprecipitated using Abcam ab13939 CENP-A antibody coupled to Dynabeads M-270 Epoxy. Chromatin extracts were incubated with antibody bound beads for 16 h at 4 °C. Bound complexes were washed once in buffer A (20 mM HEPES at pH 7.7, 20 mM KCl, 0.4 mM EDTA and 0.4 mM DTT), once in buffer A with 300 mM KCl and finally twice in buffer A with 300 mM KCl, 1 mM DTT and 0.1% Tween 20.

### DNA extraction

Following elution of the chromatin from the beads, Proteinase K (100 μg/ml) was added, and samples were incubated for 2 h at 55°C. DNA was purified from proteinase K treated samples using a DNA purification kit following the manufacturer instructions (Zymo Research, CA, USA) and was subsequently analyzed either by running a 2% low melting agarose (APEX) gel or by an Agilent 2100 Bioanalyzer by using the DNA 1000 kit.

### ChIP-Seq Library Generation and Sequencing

ChIP libraries were prepared using NEB Ultra II (Cat # E7103L) following NEBNext protocols with minor modifications. To reduce biases induced by PCR amplification of a repetitive region, libraries were prepared from 80-100 ng of input or ChIP DNA. The DNA was end-repaired and A-tailed and NEBNext adaptors (Cat # E7335S) were ligated. The libraries were PCR-amplified using only 5-7 PCR cycles since the starting DNA amount was high. Libraries were run on a 2% agarose gel and size selected for 200-350 bp. Resulting libraries were sequenced using 150 bp, paired-end sequencing on a HiSeq X instrument per manufacturer’s instructions (Illumina, San Diego, CA).

### Mapping of CENP-A-bound reads

Reads generated from PD-NC4 CENP-A ChIP-seq and from publicly available data sets were assessed for quality using FastQC (https://github.com/s-andrews/FastQC), trimmed with Sickle (https://github.com/najoshi/sickle; v1.33) to remove low-quality 5′ and 3′ end bases, and trimmed with Cutadapt (v.2.10) to remove adapters.

Processed CENP-A ChIP reads were aligned to the CHM13 whole-genome assembly v1.1 using BWA (v0.7.17) with the following parameters: bwa mem -k 50 -c 1000000 [index] [read1.fastq] [read2.fastq] for paired-end data. The resulting SAM files were filtered using SAMtools47 with FLAG score 2308 to prevent multi-mapping of reads. This filter randomly assigns reads mapping to more than one location to a single mapping location, preventing mapping biases in highly identical regions. Alignments were normalized with deepTools77 (v.3.3.0) bamCompare with the following parameters: bamCompare -b1 [ChIP.bam] -b2 [Input.bam] --operation ratio --binSize 50 -o [output.bw]. Wiggle tracks for publicly available CUT&RUN data without bulk input were generated with bamCoverage using the following parameters: bamCoverage -b [ChIP.bam] -- binSize 50 -o [output.bw].

To visualize high-confidence CENP-A positions, we used mapping scores (MAPQ: 20, or the probability of correctly mapping to another location is 0.01) to identify reads that aligned uniquely to low-frequency repeat variants. We also identified high-confidence CENP-A positions using kmer-assisted mapping (Altemose et al., 2022) and found the results to be similar to MAPQ filtering (Supplementary Fig. 4). MAPQ filtered reads were normalized to the input using bamCompare with the following normalization parameters: bamCompare -b1 [ChIP.bam] -b2 [Input.bam] --operation ratio --binSize 50 -o [output.bw] --scaleFactors 1:1. The resulting bigWig files were visualized on the UCSC Genome Browser using the CHM13 v1.1 assembly as an assembly hub.

### ChIP-seq peak calling

Significantly enriched CENP-A peaks were determined with MACS2 (v2.2.7.1) using default parameters, -g 3.03e9 and -q 0.00001. Enrichment scores were determined as the log transformed normalized value of the ratio between ChIP-seq and background, and those with a score greater than or equal to 10 were included in our study as a high-confident enrichment set. To identify high-confidence enriched CENP-A peaks that were retained between HeLa G1 and G2 cells, BED files generated from MACS2 peak calling were compared using BEDtools (v2.30.0) intersect with the following parameters: -a <G1.bed> -b <G2.bed> -u -r -f 0.50.

## Data availability

Previously published ChIP-seq and CUT&RUN data that were re-analyzed in this study can be found at the following accession numbers: GSE111381(HeLa), GSE132193 (RPE-1), GSE60951 (HuRef), and PRJNA559484 (CHM13).

## Author Contributions

M.A.M. and Y.N.-A. conceived and designed experiments and analyses and wrote the manuscript. Y.N.-A. performed ChIP-sequencing experiments. L.L. prepared sequencing libraries. M.A.M. analyzed the sequencing data. P.L. immortalized cells. K.H.M. gave analysis instructions and guidance and provided key input.

## Acknowledgments

We thank the T2T Consortium for early access to annotation, suggested mapping strategies, and assembly data. We thank Don Cleveland for critical discussion. We thank Glenis Logsdon for technical guidance and discussion and Ben Black for reagents. This work was supported by grant (R35GM142717) from the NIH to Y.N-A.

## Conflict of Interests

The authors declare no competing interests.

## Notes

### Competing Interest Statement

The authors have declared no competing interest.

