## Supplementary Figures for "Epigenetic centromere identity is precisely maintained through DNA replication but is uniquely specified among human cells"

**Supplemental figure legends**

**Supplementary Figure 1. CENP-A deposition patterns differ dramatically between hg38 centromere models and T2T centromere sequences.** Position of CENP-A enrichment in HeLa (maroon) and PD-NC4 (black) cells when mapped to the T2T (top) and hg38 (bottom) assemblies at all centromeres (excluding centromeres 1-3, presented in Fig. 1, and centromere of chromosome 19 that shares almost all α-satellite arrays with α-satellite arrays of chromosomes 1 and 5 in hg38 assembly). Scale bar, 1 Mb.

**Supplementary Figures 2-3. CENP-A binding sites differ in conventional and high-confidence data between human cell lines.** Mapping of CENP-A-bound reads (top panel of pair) and high-confidence CENP-A-bound reads (bottom panel of pair) to the T2T assembly in HeLa (maroon), PD-NC4 (black), RPE-1 (yellow), and HuRef (green) cells at all centromeres (excluding X, 5, and 9 presented in Fig. 3). Scale bar, 1 Mb.

**Supplementary Figure 4. Comparison of conventional and high-confidence CENP-A mapping strategies to the T2T assembly.** Conventional mapping, kmer-assisted mapping, and mapq filtered mapping of CENP-A-bound reads in CHM13 and PD-NC4 cells at the centromere of chromosome 2. Annotation tracks indicate positions of centromeric satellites (CenSat), higher order repeat arrays (ASat HOR), rough HOR age based on evolutionary age of HOR-haps (HOR evolutionary age), and the location of the zone of recent HOR expansion (Zone of recent expansion).

**Supplementary Figure 5. CENP-A position is maintained through DNA replication on the T2T assembly. A.** Mapping of CENP-A-bound reads in HeLa G1 (top, maroon) and G2 (bottom, dark blue) cells to the T2T assembly at all centromeres (excluding centromeres 2 and 11, presented in Fig. 4). **B.** High resolution views of CENP-A peaks depicting the position of single CENP-A nucleosomes that are maintained from G1 to G2. **C.** High resolution views of CENP-A peaks that are not maintained from G1 to G2 next to G1 peaks that are maintained into G2. Lost G1 peaks are enriched for CENP-A at a low level. Peak values are enrichment values for CENP-A over input reported by MACS2 ($p<0.00001$, ≥ 10-fold enrichment).

**Supplementary Table 1. ChIP-seq statistics for PD-NC4 CENP-A chromatin.** Read counts and alignment statistics for CENP-A ChIP-seq performed with PD-NC4 cells and two single cell derived clones. Input and CENP-A immunoprecipitation samples are shown.

**Supplementary Table 2. CENP-A-bound reads mapped to centromeric regions.** Number of CENP-A reads mapped to centromeric regions using conventional and high-confidence (MAPQ filtered) mapping techniques. The percentage of reads retained in the high-confidence dataset varies based on the size of the starting dataset and the method used to generate sequencing libraries. HeLa, PD-NC4 and HuRef data were generated with CHIP-seq, while RPE-1 data was generated with CUT&RUN.

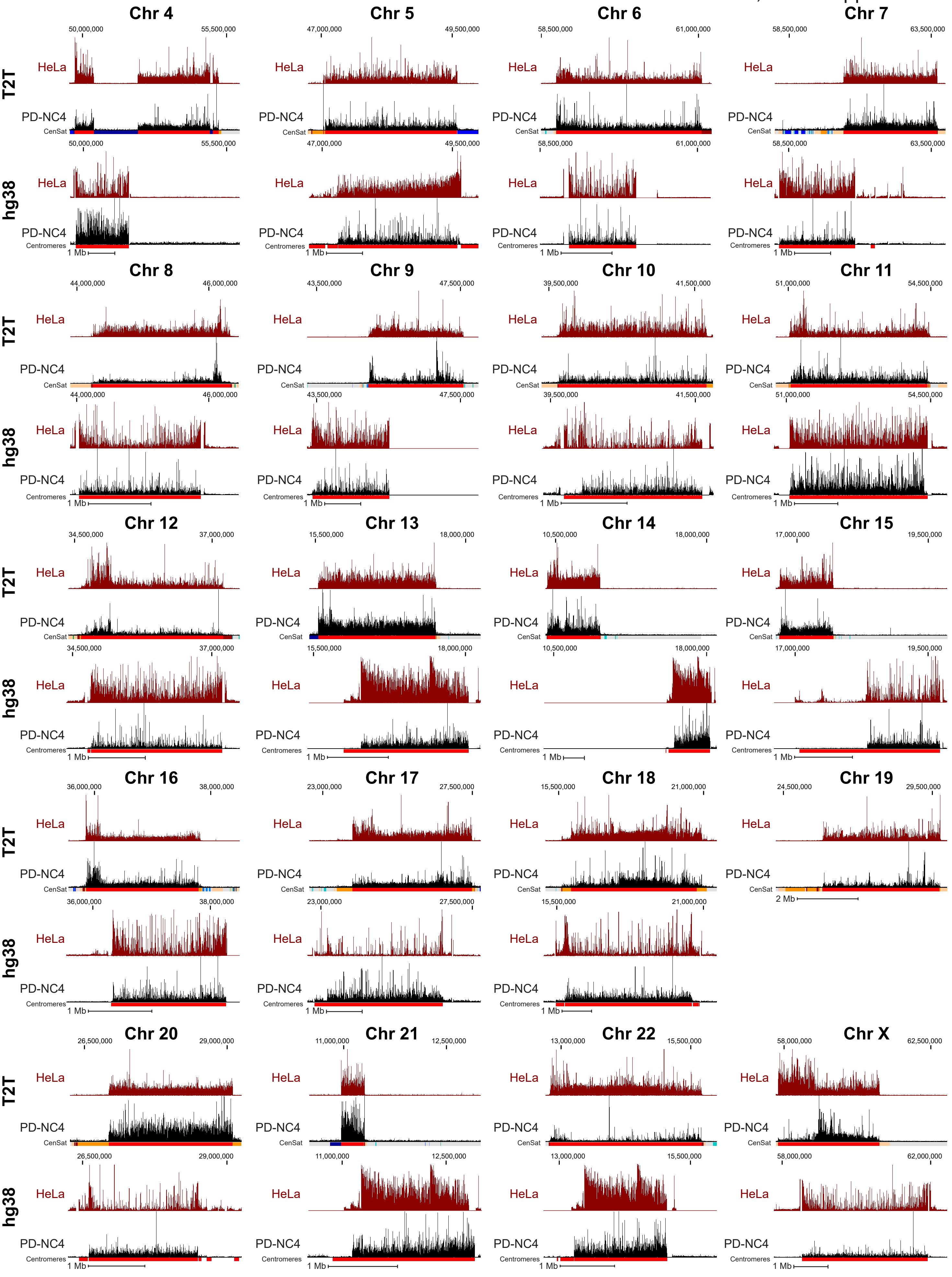


Figure S1. CENP-A deposition patterns  differ dramatically between hg38 centromere models and T2T centromere sequences.

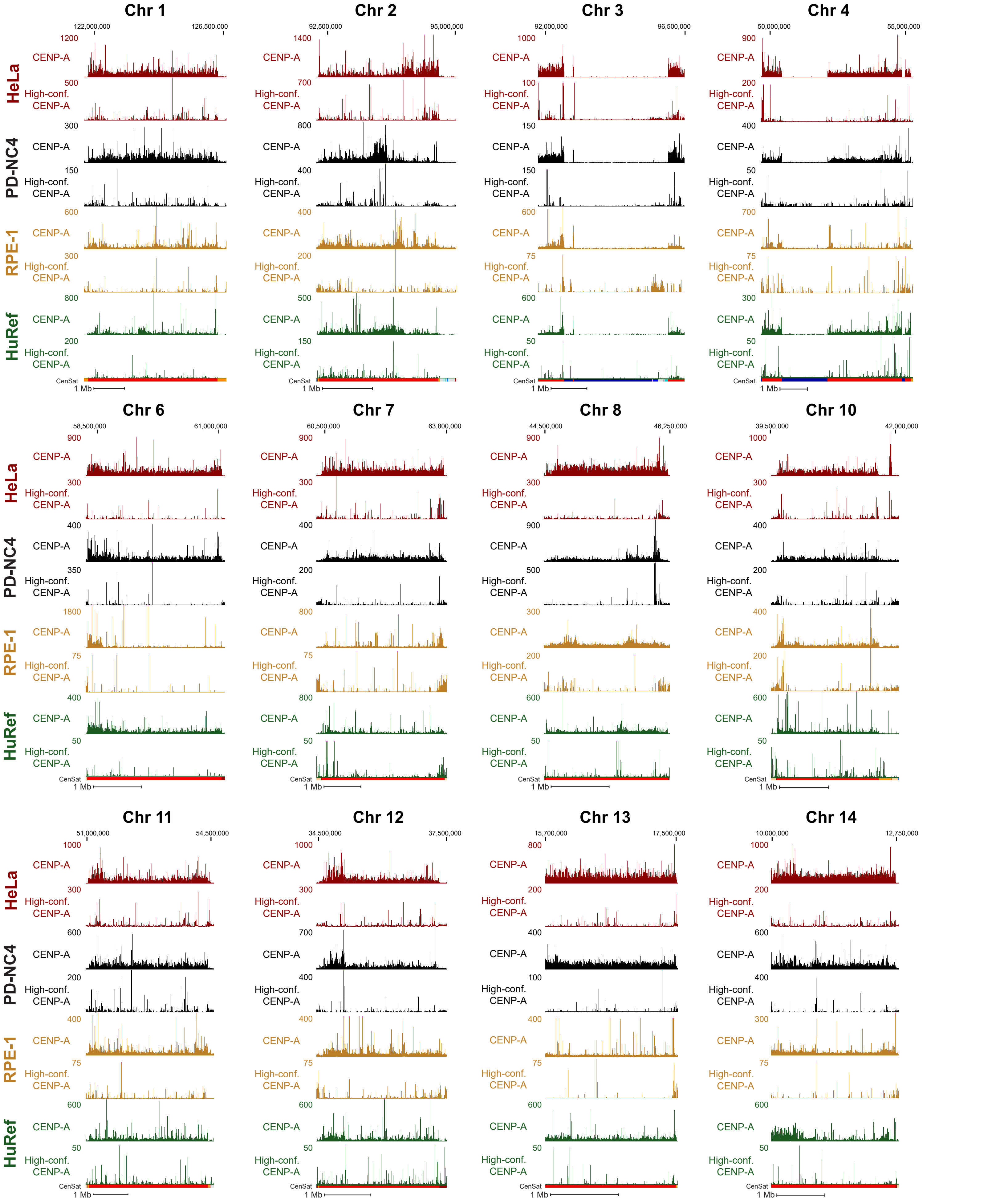


Figure S2. CENP-A binding sites differ in conventional and high-confidence data between human cell lines.

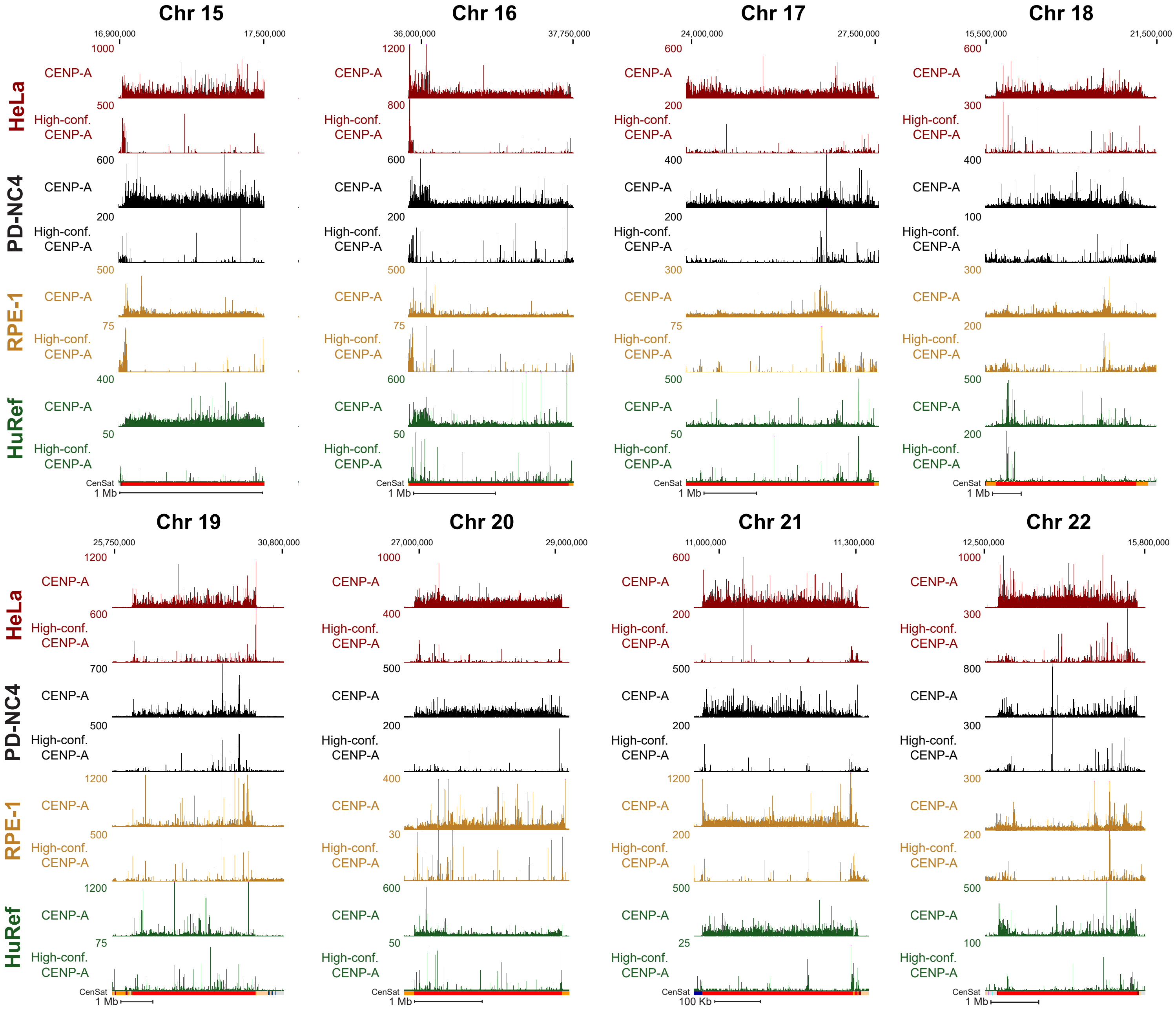


Figure S3. CENP-A binding sites differ in conventional and high-confidence data between human cell lines.

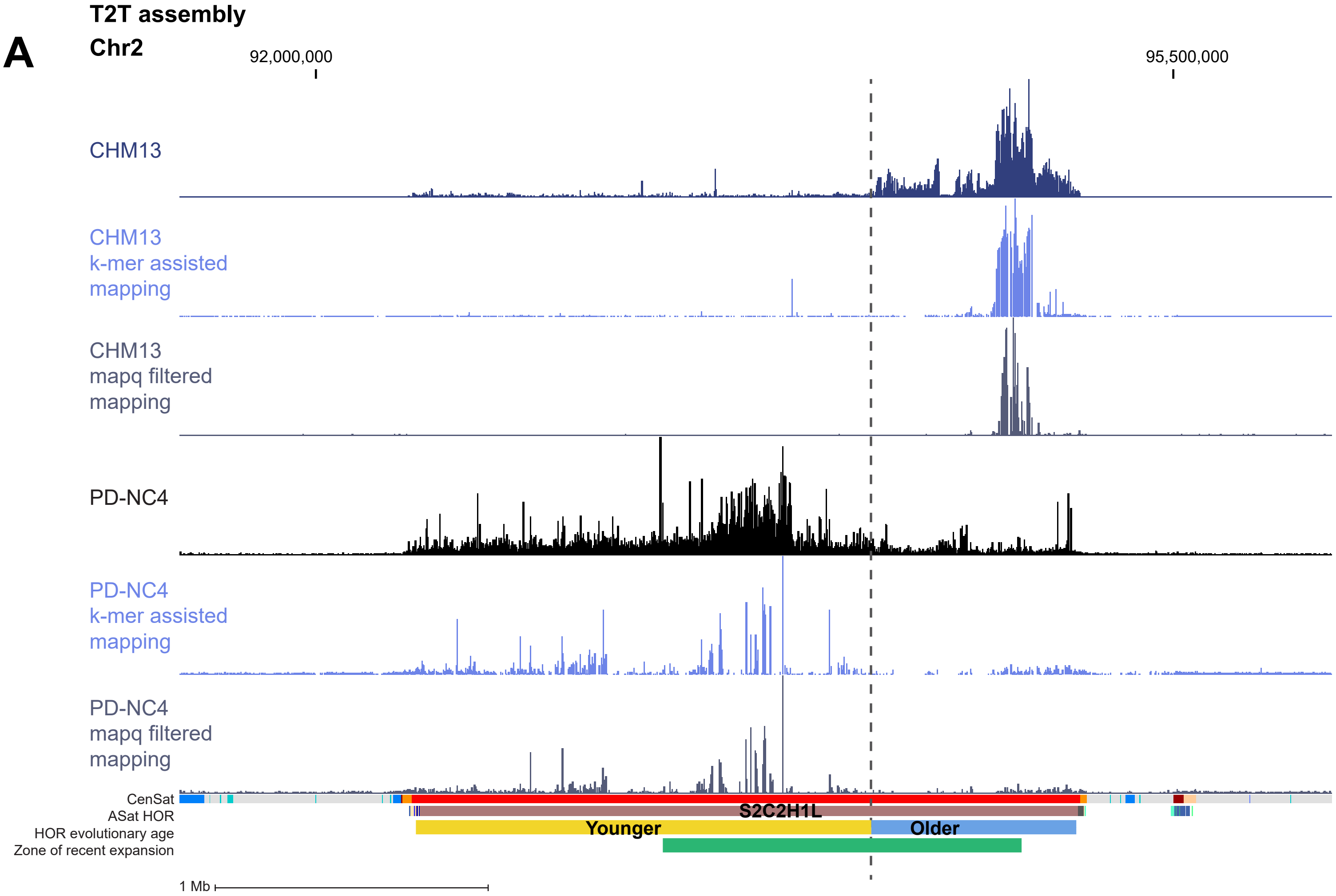


Supplemental Figure 4. Comparison of conventional and high-confidence CENP-A mapping strategies to the T2T assembly.

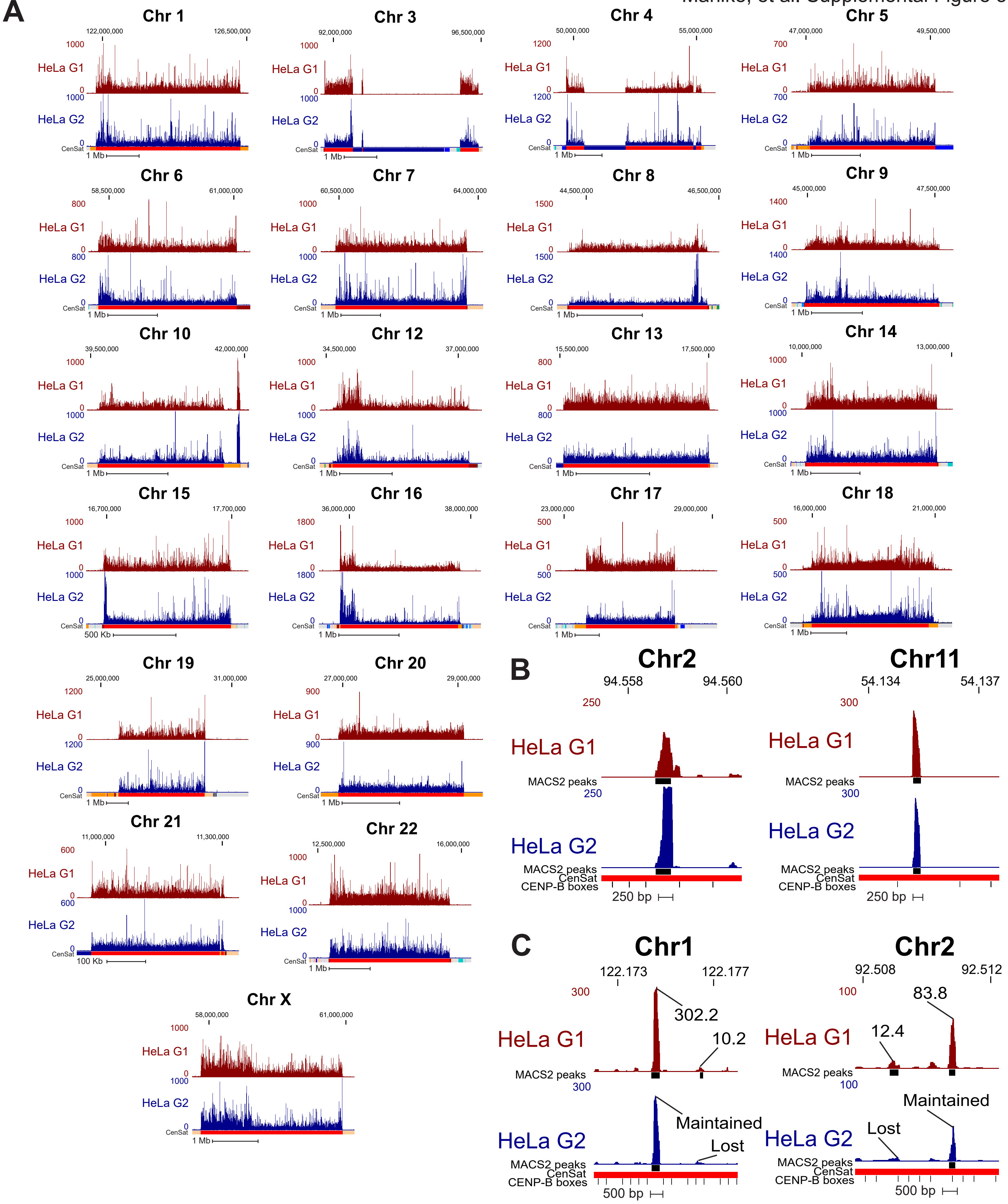


Figure S5. CENP-A position is maintained through DNA replication on the T2T assembly.

Table S1. ChIP-seq statistics for PD-NC4 CENP-A chromatin

| Type | Sample | Total number of merged paired-end reads (150bp x 2) | Total (%) number of merged reads >=100bp | Total (%) number of mapped reads |
|---|---|---|---|---|
| Input | PD-NC4 Parental | 47,895,502 | 45,566,341 (% 95.14) | 43,201,984 (% 90.20) |
| | Clone C2 | 62,196,908 | 61,245,412 (% 98.47) | 59,314,428 (% 95.37) |
| | Clone C3 | 51,806,940 | 49,831,907 (% 96.19) | 46,964,584 (% 90.65) |
| IP | PD-NC4 Parental | 30,935,042 | 30,082,060 (% 97.24) | 28,363,218 (% 91.69) |
| | Clone C2 | 38,039,172 | 35,604,664 (% 93.60) | 33,186,580 (% 87.24) |
| | Clone C3 | 34,303,265 | 32,284,577 (% 94.12) | 30,517,078 (% 88.96) |

**Supplementary Table 1. CENP-A bound reads mapped to centromeric regions.**

| Cell Line | Bulk centromeric reads | MAPQ filtered centromeric reads | % retained in MAPQ data |
|---|---|---|---|
| HeLa (G1) | 47733859 | 2688372 | 5.63 |
| PD-NC4 | 12649032 | 1306866 | 10.33 |
| RPE-1 | 5799682 | 1408048 | 24.28 |
| HuRef | 5747759 | 228340 | 3.97 |
